# Sex-specific hormone-sensitive regulatory architecture in adolescence as a scaffold for depression vulnerability

**DOI:** 10.64898/2026.05.06.723349

**Authors:** Gladi Thng, Michel Garcia-Miranda, Kailu Song, Anjali Chawla, Reine Khoury, Minh Nguyen, Gabriella Frosi, Matthew Suderman, David Liao, Natalina Salmaso, Tie Yuan Zhang, Tak Pan Wong, Yashar Zeighami, Jun Ding, Corina Nagy

**Affiliations:** McGill Group for Suicide Studies, Douglas Mental Health University Institute, Montreal, QC, Canada; Department of Human Genetics, McGill University, Montreal, QC, Canada; Quantitative Life Sciences, McGill University, Montreal, QC, Canada; Meakins-Christie Laboratories, McGill University Health Centre, Montreal, QC, Canada; Integrated Program in Neuroscience, McGill University, Montreal, QC, Canada; MRC Integrative Epidemiology Unit, University of Bristol, Bristol, UK; Population Health Sciences, Bristol Medical School, University of Bristol, Bristol, UK; Department of Neuroscience, Carleton University, Ottawa, ON, Canada; Department of Health Sciences, Carleton University, Ottawa, ON, Canada; Department of Psychiatry, McGill University, Montreal, QC, Canada; Neuroscience Division, Douglas Mental Health University Institute, Montreal, QC, Canada; Cerebral Imaging Centre, Douglas Mental Health University Institute, Montreal, QC, Canada

## Abstract

Adolescence is a critical developmental period during which sex differences in depression risk emerge, yet the underlying molecular mechanisms remain unclear. Here, we integrated longitudinal transcriptomic, epigenomic, and gene regulatory network analyses to characterize sex-specific regulatory dynamics of the brain across adolescence under normative conditions and elucidate how baseline molecular differences may shape later vulnerability. We observed that immediate early genes exhibited the strongest sex-biased signals, with males and females showing opposing developmental trajectories, suggesting a role in sex-specific stress-responsive programs. Across analyses, a consistent pattern emerged: males showed stronger activity-dependent neuronal signatures, whereas females showed stronger stress- and immune-related signatures. Extending these findings to a disease context using single-nucleus chromatin accessibility data, we identified hormone-responsive regulatory elements enriched in depression-associated regions, with sex- and cell type–specific links to genetic risk. Grounded in established models of sex-specific stress responses, these findings offer a developmental regulatory framework for understanding the differential presentation of, and vulnerability to, depression.

## INTRODUCTION

### Sex differences in typical development

Sex is a key biological variable that shapes typical development, with nearly all human complex traits exhibiting some degree of sex specificity. Interactions between genetic, hormonal, and epigenetic factors generate differential molecular landscapes in males and females^1^, as evidenced by large-scale studies reporting sex-biased gene expression and regulation across multiple human tissues^2,3^. In the healthy human brain, 9,753 genes were found to be differentially regulated in the basal ganglia^3^, and 1,239 proteins showed sex differences in expression levels in the dorsolateral prefrontal cortex^4^. These molecular differences extend to biological systems, with well-established sex differences such as in immune^5,6^ and metabolic^7^ functions that play prominent roles in the brain^8,9^. Collectively, these findings highlight the pervasive influence of sex on molecular profiles and biological functions throughout the body, including the brain.

### Sex disparities in psychiatric diseases including depression

The functional disparities in typical development can translate into differences in disease risk and manifestation, as evident by significant sex disparities in various diseases^10^. Depression is a well-established example, showing sex differences in prevalence, symptom profiles, and treatment response^11,12^. Although our understanding of sex differences in depression remains incomplete, numerous studies have reported evidence for sex-specific mechanisms at multiple biological levels, including transcriptional pathways^13,14^, as well as structural and molecular differences in the brain^15^. Notably, recent single-nucleus RNA-sequencing studies of postmortem human brain tissues reported that distinct cell populations are differentially affected in depressed males versus females^16,17^. These findings highlight functional sex differences in depression and underscore the need for a clearer understanding of sex-specific mechanisms to better elucidate its biological basis.

### Adolescence and puberty as a critical period

Although sex differences are present throughout life, adolescence, and specifically puberty, represent a period when sex-biased developmental processes are most pronounced. This is evident in postmortem human brain data identifying over 2000 genes with sex-biased expression at each developmental stage from prenatal to adulthood, with peak differentiation occurring at puberty with 4164 genes^18^. Gonadal hormones exert both organizational and activational effects on the brain, with organizational effects producing lasting structural and functional changes during sensitive developmental windows, and activational effects that “activate” neural tissues previously organized by sex, reflect transient, reversible modulation^19,20^. Puberty represents a second such sensitive window, following the prenatal/early postnatal period, during which hormonal surges may contribute to lasting changes in cellular hormone sensitivity, and shape transcriptomic and epigenomic landscapes in the brain^21,22^. Notably, this critical window coincides with the emergence of many mental disorders, including depression, whose prevalence rises to approximately twice the rate in females compared to males^23,24^. These organizational and activational effects of pubertal hormones may therefore contribute to this disparity by establishing divergent baselines in stress-sensitive circuits^25–27^—changes that, given the persistent female preponderance in depression risk across the lifespan, may have consequences extending well beyond adolescence.

### Longitudinal profiling to assess depression risk

The findings so far suggest that puberty is a period of dynamic gene expression and neural remodeling, and these developmental changes differ between sexes. Sex-specific expression trajectories during this window may therefore reveal baseline molecular patterns associated with depression-related genes. However, most existing studies either focus on post-onset cases, examine gene expression without considering regulatory context, or do not explicitly account for sex differences. Consequently, the normative developmental trajectories through which sex-biased transcriptional programs emerge remain insufficiently characterized. Profiling these trajectories prior to the emergence of disease phenotypes provides an opportunity to define baseline molecular programs that may be relevant to later depression risk. To address these gaps, we investigated normative gene regulatory dynamics using a pseudo-longitudinal study of the mouse prelimbic cortex (an analogue of the human prefrontal cortex), focusing on sex-specific trajectories of depression-associated genes. This framework enabled characterization of the emergence and evolution of sex-biased molecular patterns across development, providing insight into the regulatory and transcriptional landscapes that may contribute to sex differences in depression vulnerability.

## RESULTS

Our transcriptomic dataset comprised 20 high-frequency timepoints spanning adolescence (P28 to P72), along with one distal adult timepoint at P120^28^ (Figure 1A). Time Point Selection analysis^29^ identified five of these timepoints (P28, P32, P42, P48, P72) as the most biologically informative (Figure S1), and a complementary DNA methylation dataset comprising these five timepoints was analyzed in parallel. Given the use of bulk tissue, cell type deconvolution was performed for both datasets to assess potential variation in cellular composition (Figure S2-3). No sex- or age-associated differences in estimated cell proportions were detected in the transcriptomic data, while only age-related shifts were observed in the methylation data and accounted for in downstream analyses.

**Figure 1.**
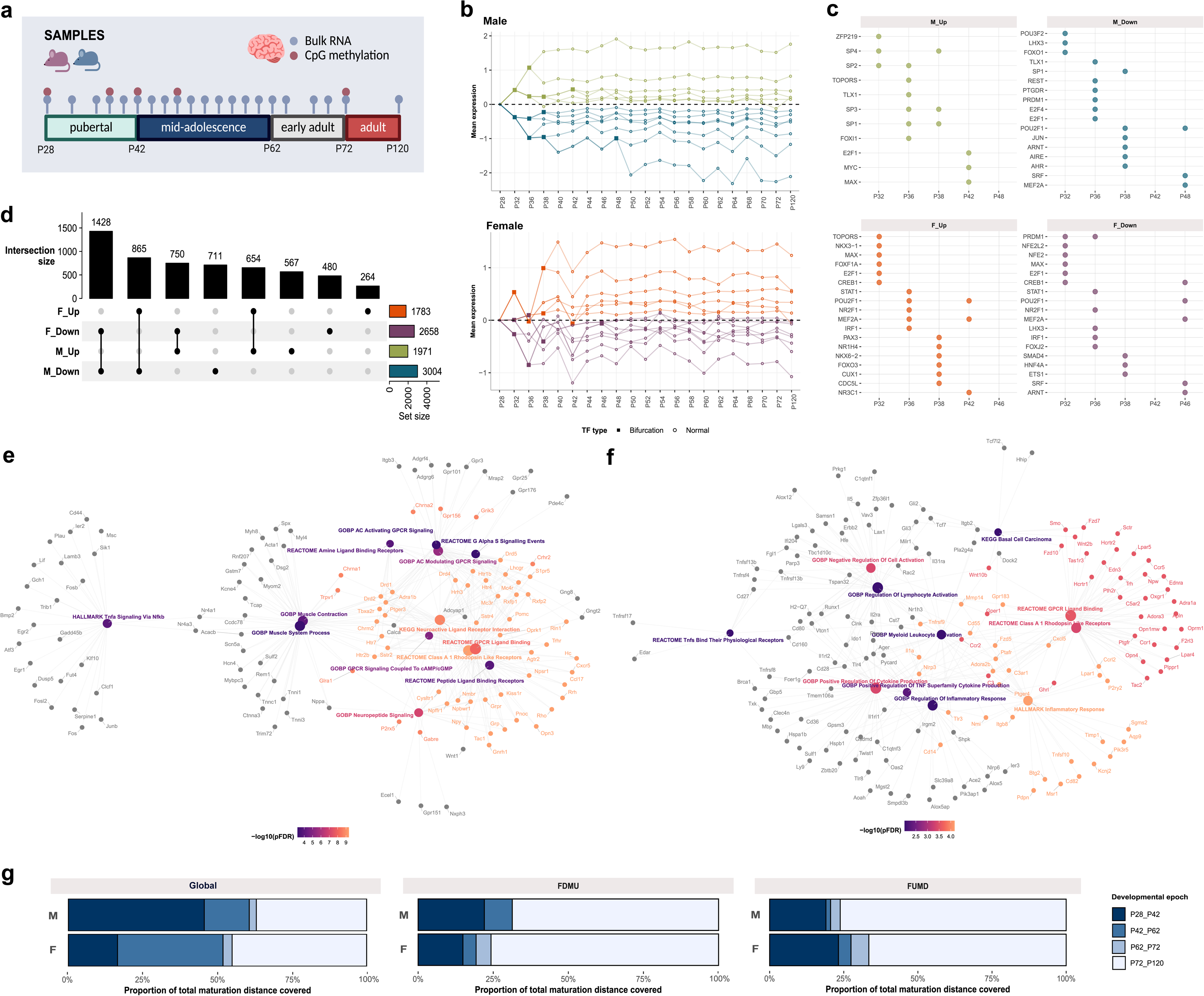
Global sex-divergent developmental regulatory trajectories prioritize neuromodulatory signaling in males and immune signaling in females. (A) Graphical overview of study design. (B) Male-and female-specific trajectories constructed using IDREM. Up- and downregulated paths relative to baseline are shown in different colours for each sex. Bifurcation nodes, representing points at which a given path branches into a new path, are represented as squares. (C) The top 3 initiator TFs at each bifurcation node for each path category. (D) Network consisting of significant pathways for FDMU genes and (E) FUMD genes. (F) Distance to adulthood (P120) when using the top variable genes from the whole transcriptome (left), all FDMU genes (center) and all FUMD genes (right) for each sex.

### Global sex-divergent developmental regulatory trajectories prioritize neuromodulatory signaling in males and immune signaling in females

To characterize broad developmental trajectories, we constructed dynamic gene regulatory networks across all timepoints using iDREM^30^, which assigns genes to temporal expression paths and infers candidate upstream regulators of each trajectory, hereon referred to as networks. Regulators (hereafter TFs) include canonical transcription factors and model-inferred upstream regulators capturing direct and indirect regulatory effects. Networks were generated separately for females and males to capture sex-specific regulatory dynamics. Relative to baseline P28, females exhibited 6 upregulated and 8 downregulated trajectories over development, whereas males showed 6 upregulated and 7 downregulated trajectories (Figure 1B; Figure S4).

We next quantified overlap in gene membership across trajectories between sexes to identify concordant and divergent regulation of gene programs. Genes downregulated in both sexes showed the largest overlap, consistent with shared developmental refinement processes. The next largest overlaps, however, comprised genes regulated in opposite directions between sexes (Figure 1D), indicating substantial divergence in developmental prioritization. Pathway enrichment analysis revealed that genes downregulated in females but upregulated in males (FDMU) were significantly enriched for G-protein–coupled receptor signaling (pFDR<0.001), with neuroactive ligand–receptor interaction as the top term (pFDR<0.0001) (Figure 1E). This set included dopaminergic, serotonergic histaminergic, glutamatergic, and peptidergic receptors. In contrast, genes upregulated in females but downregulated in males (FUMD) were mainly enriched for immune-related processes (pFDR<0.0001), including cytokine production, lymphocyte activation, and most significantly the Hallmark inflammatory response pathway (Figure 1F).

To identify regulatory factors underlying these pathways, we examined TFs associated with trajectory bifurcation (initiator TFs; Figure 1C), which occurred primarily in the earlier timepoints between P32 and P46. While there are master TFs present in multiple categories (e.g., *Pou2f1*, *Stat1*, *E2f1*, *Mef2a*, *Max*), there are notable unique TFs in each category that distinguish sex- and direction-specific programs. For instance, male upregulated trajectories included the *Sp* family (*Sp1–4*; neuronal regulation), while downregulated trajectories featured *Rest* (neuronal gene repressor in non-neuronal cells) and *Jun* (immediate early gene). In females, upregulated trajectories highlighted *Nr3c1* (glucocorticoid receptor) and *Nkx6-2* (oligodendrocyte differentiation), pointing to stress-responsive and glial-associated regulatory programs, while downregulated trajectories featured *Nfe2l2* (redox/stress response) and *Hnf4a* (master metabolic regulator). Across the broader TF set (Figure S5), a consistent pattern emerged regardless of regulatory direction: early trajectories were driven by largely sex-distinct initiator TFs, whereas maintenance TFs at P120 showed greater overlap, suggesting convergence in regulatory architecture by adulthood.

To contextualize the sex-biased pathway-level differences in overall brain development, we quantified transcriptomic maturation revealing distinct trajectories between sexes (Figure 1G). Globally, males showed rapid early-adolescent progression, covering 46% of the total developmental distance between P28 and P42. Females progressed more slowly (∼17% between P28 and P42), with larger shifts occurring from P42 to P62 (35%) onwards. Both sexes showed minimal progression between P62 and P72, followed by substantial progression from P72 to P120. These differences were more pronounced within sex-biased gene sets. For FDMU genes, males exhibited greater early progression (22% between P28 and P42) than females (15%), whereas this pattern was reversed for FUMD genes, with females showing greater early progression (23%) relative to males (19%). Together, these findings indicate that transcriptomic maturation is sex-specific and dependent on the underlying molecular systems being prioritized.

### Sex-biased depression-associated genes converge on an activity-dependent transcriptional module integrating stress, immune, and neuronal signaling

As the iDREM networks were constructed separately for females and males precluding statistical comparison, we next identified genes with sex-divergent developmental dynamics by performing differential expression analysis across autosomal and X-linked genes using non-linear generalized additive models (GAMs). Genes were classified as differentially expressed (DEGs) if they differed between sexes in overall expression, developmental trajectory, or both. At pFDR<0.05, 143 DEGs were identified, with 82 genes showing higher overall expression in males (male-biased) and 61 female-biased genes (Figure 2A; Table S1). To assess enrichment for depression-associated genes, we compiled a curated gene list (Table S2). Of the 143 DEGs, 47 were depression-associated (depDEGs; bold highlight), representing a significant enrichment relative to expectation (p=2×10^-4^) and suggesting a molecular vulnerability induced at this developmental transition. This enrichment was observed in both sexes, with 26/82 male-biased DEGs and 21/61 female-biased DEGs identified as depDEGs. The proportion of depression-associated genes did not differ significantly between sexes (p=0.857), suggesting a balanced molecular vulnerability.

**Figure 2.**
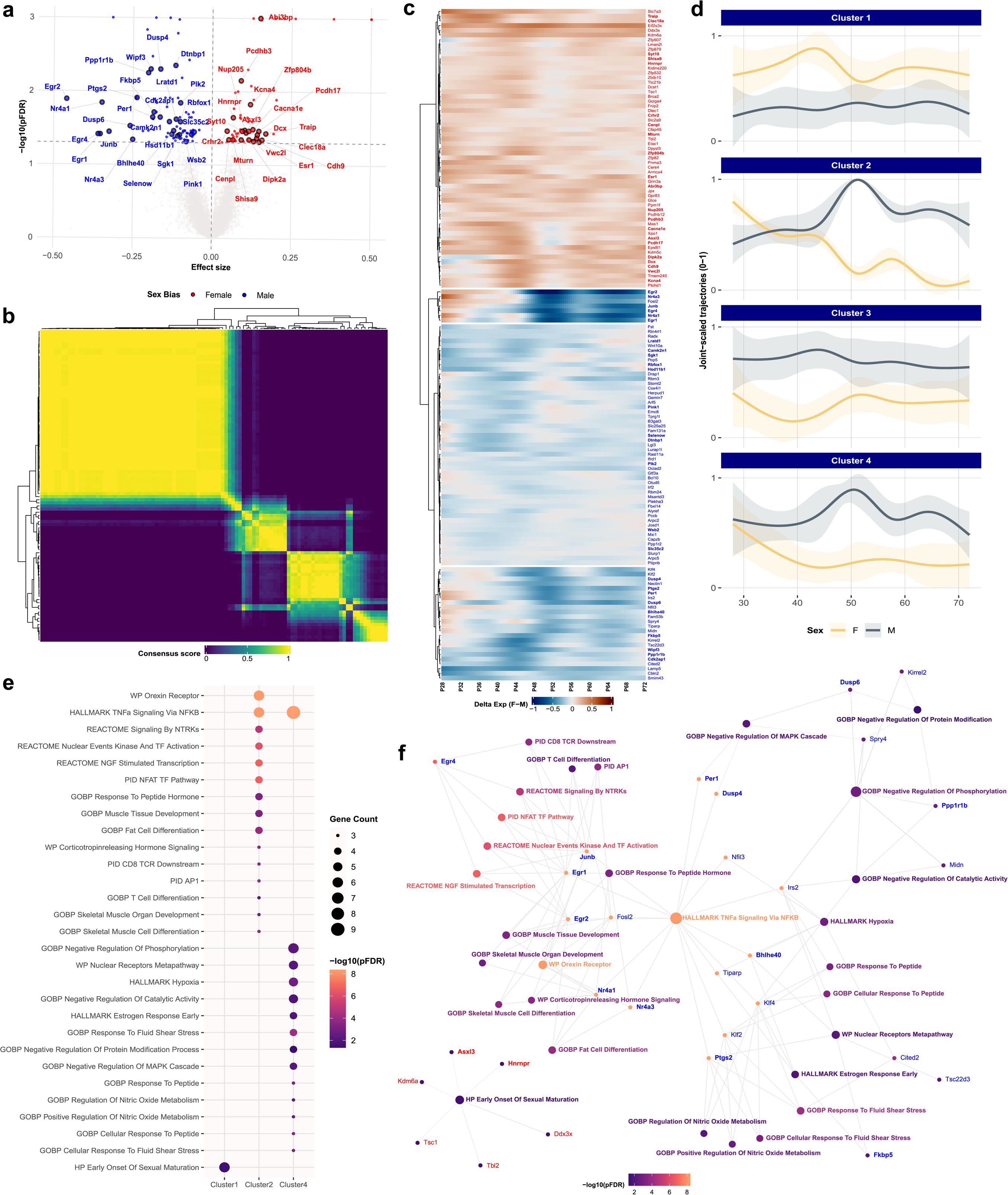
Sex-biased depression-associated genes converge on an activity-dependent transcriptional module integrating stress, immune, and neuronal signaling. (A) Volcano plot showing all identified sex DEGs, with female-biased DEGs in red and male-biased DEGs in blue. Depression-associated DEGs (depDEGs) are highlighted with a black border and the top 50 depDEGs are labelled. Effect sizes correspond to the estimated coefficients of the linear sex term in the GAM framework, as differential expression was primarily driven by this term. (B) Heatmap showing the consensus score of each gene upon running multiple iterations of hierarchical clustering of delta expression trajectories (female minus male). (C) Heatmap showing the delta expression trajectories, grouped into 4 clusters. Female-biased DEGs are in red, male-biased DEGs are in blue, and depDEGs are in bold. *Xist* and *Tsix* were excluded from the heatmap due to their large bias towards females that would skew the heatmap. (d) The average delta-expression trajectory for each cluster, with the shaded area representing the confidence interval. (E) Pathways that were significantly enriched in each cluster of DEGs at pFDR<0.05. No significant enrichment was observed for cluster 3. (F) A combined network showing all significantly enriched pathways across clusters and the DEGs involved.

To characterize how these sex differences unfold across adolescence, expression trajectories were jointly scaled and clustered. This revealed four distinct trajectory patterns (Figure 2B), with most DEGs maintaining consistently higher expression in one sex, supporting the GAM results (Figure 2C-D). Notably, cluster 2, enriched for immediate early genes (IEGs, e.g., *Junb*, *Egr1/2/4, Nr4a1/3*), showed higher expression in females early in development, followed by markedly higher expression in males later. Cluster 4, enriched for stress-responsive genes (e.g., *Per1*, *Bhlhe40*, *Dusp4/6*, *Fkbp5*, *Tsc22d3*), displayed a similar temporal pattern, although without a complete crossover in trajectory between sexes. Pathway enrichment of each cluster revealed biologically coherent enrichments for clusters 1, 2, and 4 (pFDR<0.05; Figure 2E). Cluster 1 comprised female-biased chrX genes enriched for processes involved in early sexual maturation. Cluster 2 was highly enriched for Hallmark TNF-α signaling via NF-κB along with several signaling pathways relevant to neuronal function, such as orexin receptor pathway (arousal and modulation of cortical plasticity), NFAT signaling (neuronal excitability and plasticity), NGF-stimulated transcription (dendritic growth, synaptic maturation). Cluster 4 was likewise highly enriched for Hallmark TNF-α signaling via NF-κB, along with Hallmark estrogen response early, Hallmark hypoxia, and nuclear receptors metapathways (sensing of steroid/thyroid hormones to regulate expression). Collectively, these enrichments suggest substantial convergence between inflammatory, hormonal, stress-responsive, and neuronal signaling programs within a shared regulatory framework (Figure 2F).

### Sex-biased promoter DNA methylation of stress-, immune-, and neuronal X-linked genes during adolescence is associated with loci containing steroid hormone receptor binding motifs

To determine whether sex-biased gene expression is associated with DNA methylation differences, we analyzed a complementary methylation dataset using the same GAM framework. We identified 6333 differentially methylated CpGs (DMPs) exhibiting baseline sex differences (Table S3). Of these, 4295 were hypermethylated in females and 2038 were hypermethylated in males, indicating a global excess of female-biased hypermethylation across chromosomes (Figure 3A). On autosomes, promoter-associated hypermethylation were more frequent in males (35%) than females (23%; Figure 3B), whereas the pattern was reversed for chrX (34% in females vs 12% in males), consistent with X-chromosome inactivation (XCI) and dosage compensation.

**Figure 3.**
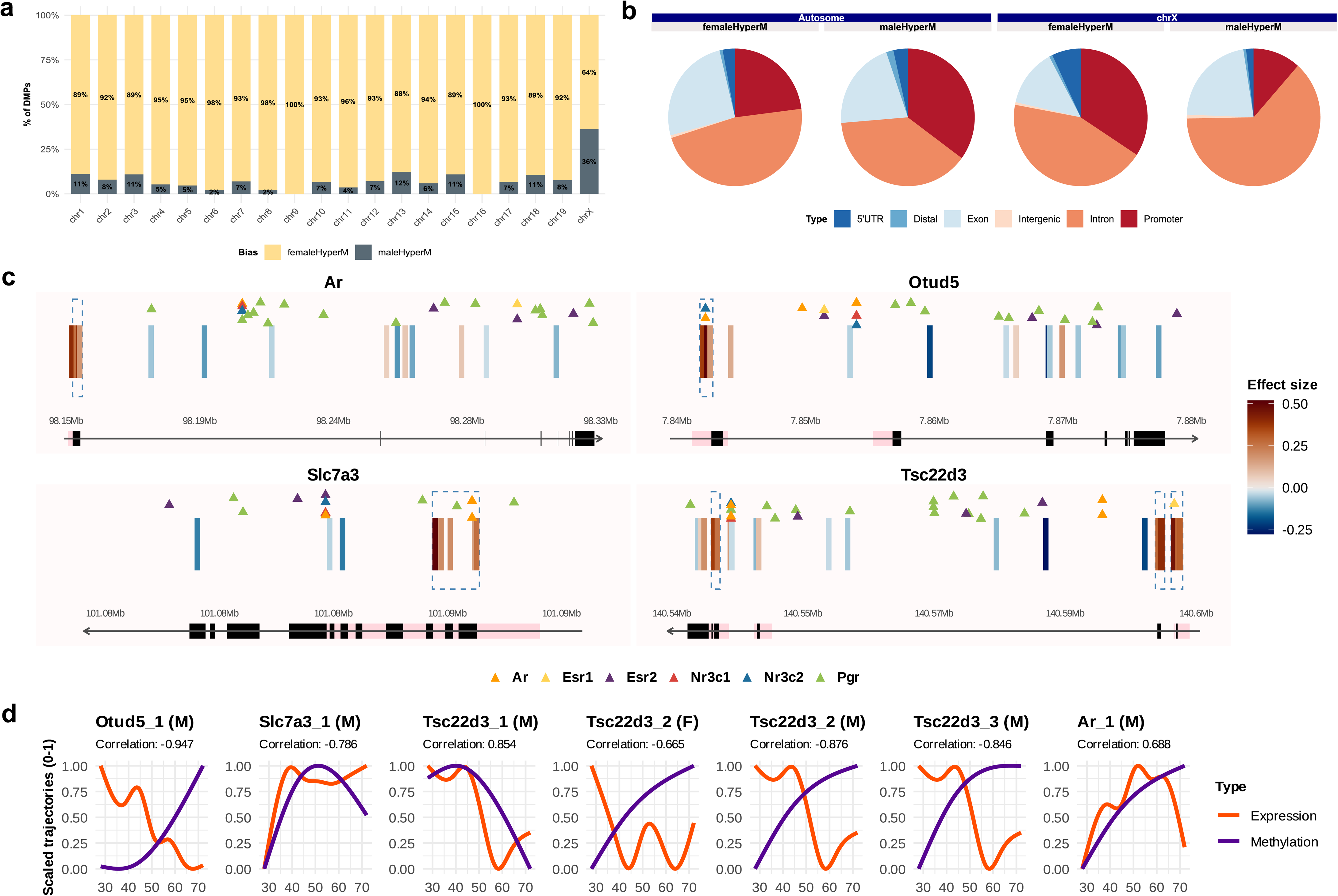
Sex-biased promoter DNA methylation of stress-, immune-, and neuronal X-linked genes during adolescence is associated with loci containing steroid hormone receptor binding motifs. (A) Barplot showing the distribution of hypermethylated DMPs per chromosome, coloured by sex bias. (B) Pie charts showing the genomic context of DMPs located in autosomes and chromosome X separately. (C) Gene tracks showing the location of each DMP (each individual bar) per gene, coloured by sex bias. Along the gene track, promoters are in pink and exons in black. DMRs are highlighted in a dotted blue box, and hormone motifs are represented as triangles, colored by hormone. Only genes with identified DMRs are shown. (D) Correlations between the average methylation trajectory of a given DMR and the expression trajectory. Only significant correlations are shown (p<0.05).

Across autosomes and chrX, 109 DMPs mapped to 17 sex DEGs. To focus on coordinated regulatory changes, we identified putative differentially methylated regions (DMRs) by clustering nearby DMPs with the same direction of methylation change (Figure S6). For chrX DEGs with DMRs, we focused on those previously classified as non-escapees of XCI^31,32^, as the presence of sex-biased expression in these genes that are expected to show dosage equivalence between sexes suggests additional regulatory modulation potentially involving DNA methylation. This yielded promoter-associated DMRs in *Tsc22d3* (a mediator of glucocorticoid-induced immunosuppression; contained 3 DMRs), *Slc7a3* (a cationic transporter linked to neuromodulatory amino acid availability), *Otud5* (a regulator of innate inflammatory pathways) and *Ar*, all hypermethylated in females (Figure 3C). Although *Ar* was not identified as a DMP, it was retained due to the high density of associated DMPs and its functional relevance as a nuclear hormone receptor. When intersecting these promoter DMRs with steroid hormone receptor binding motifs (*Ar*, *Esr1/2*, *Nr3c1/2*, *Pgr*), *Slc7a3* DMR overlapped *Ar* and *Pgr* motifs, *Tsc22d3* DMR #3 overlapped *Esr1* motif, and *Otud5* DMR overlapped *Ar* and *Nr3c2* motifs (p-value cutoff: 0.0001; Figure 3D; Table S4), suggesting that epigenetic remodeling at these loci may occur within hormone-responsive regulatory contexts.

To determine the functional regulatory relevance of these promoter DMRs, we next tested whether methylation changes at these DMRs tracked expression across development by correlating the fitted methylation trajectory of each DMR with its corresponding gene expression trajectory in a sex-specific manner. Both temporal similarity and directional association were quantified using distance and Spearman correlations, respectively (see Methods). Significant correlations were observed for all DMRs in males and only for *Tsc22d3* (DMR #2) in females (p<0.05; Table S5), with promoter hypermethylation generally associated with reduced expression. The predominance of significant correlations in males is consistent with allelic masking by XCI in females, where methylation dynamics on the active allele may be obscured by the inactive allele. Notably, *Tsc22d3* DMR #2 showed correlated trajectories in both sexes, suggesting that methylation-coupled regulation at this glucocorticoid-responsive locus is robust to dosage compensation effect. Together, these findings support a model in which hormone-sensitive promoter methylation is associated with sex-specific expression of stress- and immune-related X-linked genes during adolescent brain development.

### Sex-biased DEGs in stress, immune and neuronal signaling programs show differential regulatory targeting across adolescence

As disentangling XCI-driven effects from independent methylation regulation remains challenging, the extent to which DNA methylation independently contributes to sex-biased expression is difficult to fully resolve. Moreover, methylation analyses primarily capture local chromatin state at individual loci and may not fully reflect broader differences in upstream regulatory architecture. Consistent with this, iDREM identified distinct TFs regulating overlapping gene sets in males and females, suggesting that sex-divergent transcription may also arise through differences in TF targeting. We therefore extended our analysis to sample-specific gene regulatory networks (GRNs) using LIONESS^33^ to identify where sex differences in transcriptional regulation emerge during adolescent brain development. LIONESS edge weights quantify the inferred strength of a regulator–target relationship, with higher positive values indicating stronger evidence of regulatory influence and negative values indicating little or no evidence of regulatory influence.

Applying a GAM framework, 86,204 sex-biased edges comprising approximately 10% of all edges were identified at pFDR<0.05. Of these, 30,565 showed higher evidence of regulation in females and 55,639 in males, with differences in overall regulatory strength rather than pronounced non-linear developmental changes. Among these, 1134 sex-biased edges connected 288 TFs to 44 DEGs of which, 15 are depDEGs (Table S6; Figure S7). To test regulatory-expression coupling, we correlated the fitted expression trajectory of each DEG with all regulatory trajectories involving that gene in a sex-specific manner (Figure S7; Table S8). As LIONESS edge weights are not inherently directional, the sign of the correlation reflects whether regulatory strength is positively or negatively associated with gene expression, consistent with putative activating or repressing relationships. In males, distance and Spearman correlation p-values were highly concordant (r=0.94, p<0.0001) with similar effect size patterns. In females, concordance was weaker (r=0.64, p<0.0001) and Spearman correlation was generally lower than distance correlations (Figure 4A). This pattern suggests that the regulation-expression relationships are largely monotonic in males but involve more complex non-linear dynamics in females. At p<0.1, the remaining correlations involving 36 DEGs (12 depDEGs) and 256 TFs showed strong effect sizes (|r|>0.5; Figure 4B), indicating robust regulatory-expression coupling in both sexes.

**Figure 4.**
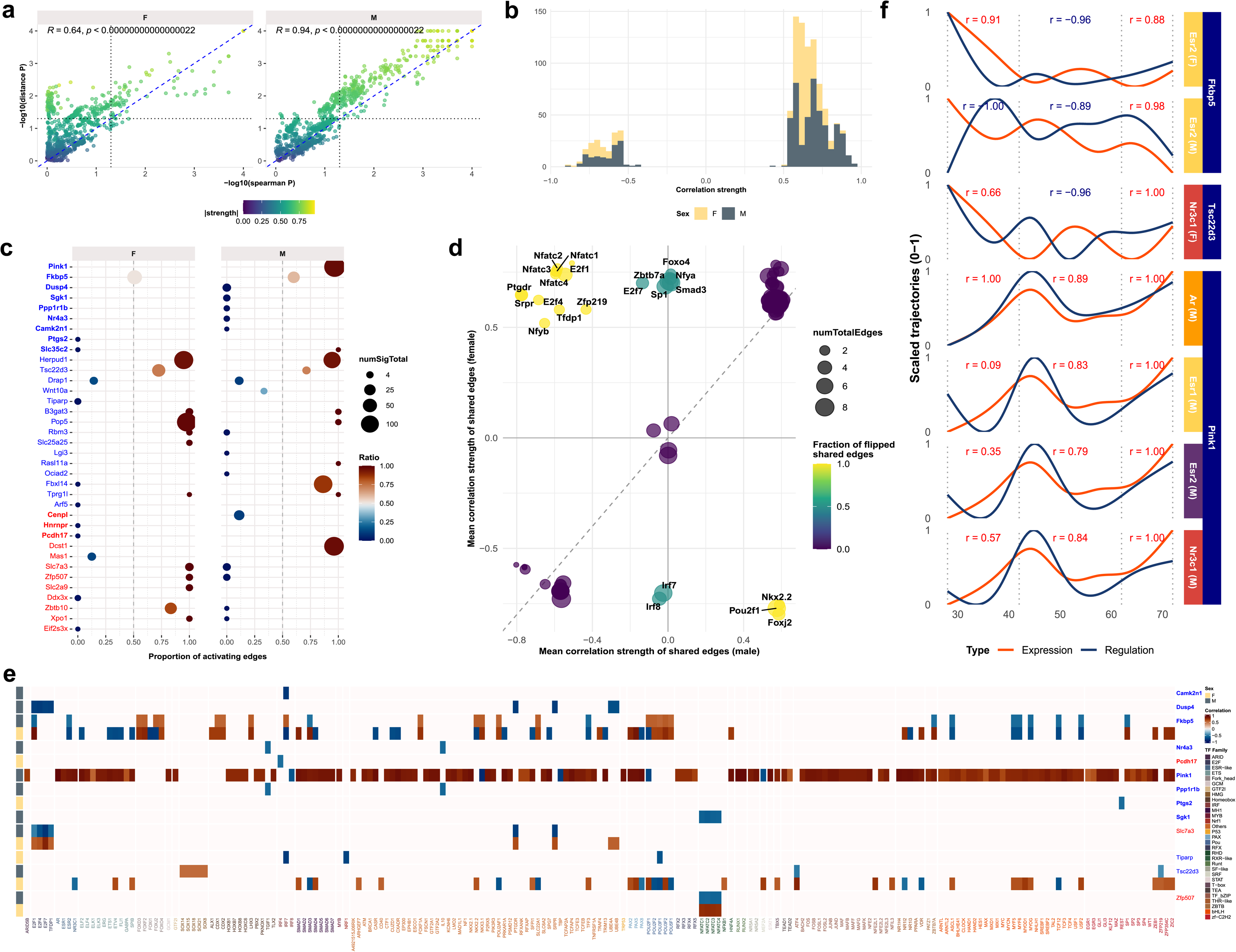
Sex-biased DEGs in stress, immune and neuronal signaling programs show differential regulatory targeting across adolescence. (A) Scatterplot showing the correlation between p-values for distance correlation and Spearman correlation for males and females separately. (B) Histogram showing the distribution of absolute hybrid correlation values after filtering for correlations with p<0.1. (C) Summary plot of DEGs showing the proportion of putatively activating regulatory edges relative to total regulatory edges per gene. 0 represents pure repression while 1 represents pure activation. (D) Summary plot of TF-specific regulatory consistency across shared TF–target edges between sexes, including direction concordance and average correlation strength. (E) Heatmap to show the specific TFs involved in regulating each DEG, coloured by activating (red) or repressive (blue) upstream control. Only selected DEGs are shown in this plot. (F) Regulation and expression trajectories of DEGs that have significant regulation-expression coupling with hormone-associated regulatory signals.

To better distil these results, we derived a summary measure per DEG per sex to quantify the relative strength and direction of regulatory input (Figure 4C). Genes were considered more strongly regulated when TFs exhibited greater concordance in regulatory direction (indicated by darker color) and/or when a larger number of TFs targeted the gene (indicated by circle diameter). Genes involved in activity- and stress-responsive signaling relevant to neuronal plasticity (e.g., *Camk2n1*, *Dusp4*, *Nr4a3*, *Ppp1r1b*, *Sgk1*, *Pink1*) showed stronger regulatory input in males. Regulatory associations for most of these genes were predominantly negative in males, suggesting a bias toward repressive upstream control, with the exception of *Pink1*, which displayed activating regulatory input. In contrast, genes associated with glucocorticoid-responsive, inflammatory, and cellular homeostatic processes (e.g., *Fkbp5*, *Tsc22d3*, *Pcdh17*, *Tiparp*, *Ptgs2*, *Hnrnpr*, *Slc25a25*, *Slc2a9*) showed stronger regulatory input in females. Several genes displayed opposing regulatory biases between sexes, such as *Slc7a3*, *Zfp507*, *Zbtb10*, and *Xpo1*, which were predominantly repressed in males but activated in females, indicating sex-specific directionality of upstream control. Notably, *Fkbp5* and *Tsc22d3* showed comparatively tight regulation in both sexes (closer proximity to the midline), particularly in females, reflecting a more balanced contribution of activating and repressing inputs. Examination of upstream regulators (Figure 4E) further revealed that *Fkbp5* was largely regulated by concordant TF inputs in both sexes, whereas *Tsc22d3* exhibited stronger and more diverse regulatory input in females, while in males it was primarily influenced by activating *Sox* family TFs. In males, *Camk2n1*, *Dusp4*, *Nr4a3*, *Ppp1r1b*, and *Sgk1* were predominantly targeted by immune- and stress-related TFs (e.g., *Irf7*, *Ptgdr*, *Il10*, *Nfatc1–4*). In contrast, genes such as *Slc7a3* and *Zfp507* exhibited consistent sex-specific differences in regulatory directionality across multiple TFs (e.g., *E2f* family, *Ptgdr*, *Srpr*).

As directionally flipped edges represent the strongest form of regulatory asymmetry, for each TF we quantified the fraction of shared significant edges that were oppositely regulated between sexes using all significant edges (Figure 4D). Members of the Nfatc family (*Nfatc1—4*), inflammation-related TFs (*Ptgdr*, *Srpr*) and cell cycle TFs (*E2f4*, *Tfdp1*, *Zfp219*) exhibited a high proportion of flipped edges with stronger regulatory-expression coupling in females. In contrast, *Nkx2-2*, *Pou2f1* and *Foxj2* showed stronger coupling in males. Intermediate patterns were observed for TFs such as *Irf7*, *Irf8*, *Zbtb7a*, *Sp1*, *Nfya*, *Smad3*, and *Foxo4*. Finally, we asked whether hormone-associated regulatory interactions show developmental modulation, focusing on genes that exhibited significant regulatory coupling with hormone receptors, including *Fkbp5*, *Tsc22d3*, and *Pink1*. As global correlation does not capture transient changes across development, we evaluated expression–regulation relationships across discrete developmental windows (P28–P42, P42–P62, and P62–P72). This revealed that hormone-associated TFs, particularly *Esr1–Fkbp5* and *Nr3c1–Tsc22d3* in females, showed consistently aligned regulatory–expression dynamics across transitions, indicating sustained hormone-linked regulatory control during adolescent development (Figure 4F). Together, these findings suggest that sex-biased transcriptional programs in stress-, immune–and neuronal-associated genes are shaped by sex-specific TF–target regulatory coupling across adolescence.

### Hormone-responsive regulatory regions in depression show cell-type–specific motif enrichment and associate with sex-specific genetic vulnerability

Across our analyses of DNA methylation and TF targeting, we observed consistent evidence that hormone-related mechanisms contribute to the regulation of DEGs and depDEGs across adolescence, in line with their established role in puberty and depression. These findings raise the possibility that hormone-linked regulatory programs established during development may persist into adulthood and contribute to disease-relevant regulatory architecture. To evaluate this in a human disease context, we leveraged cell type–resolved open chromatin regions (OCRs) from the postmortem dorsolateral prefrontal cortex of depressed and control subjects^34^. Hormone-responsive OCRs (hOCRs) were identified in both baseline OCRs, representing normative regulatory architecture, and depression-associated OCRs, reflecting disease-related chromatin remodeling (Table S8). *Esr1/2* motifs accounted for the highest percentage of hOCRs across cell types (Figure 5A). Relative to baseline, the proportion of depression-associated hOCRs generally increased across multiple cell types, especially within regions showing increased accessibility (MDD_Increase), including astrocytes, excitatory neurons and oligodendrocytes. In examining the genomic contexts of hOCRs, most cell types showed a promoter-dominant profile for baseline OCRs but not for depression-associated OCRs, indicating a shift away from proximal regulation (Figure 5B). Notably, neuronal populations exhibited a higher proportion of distal hOCRs in MDD_Increase compared to other cell types, suggesting greater engagement of enhancer-mediated regulation in neurons in depression.

**Figure 5.**
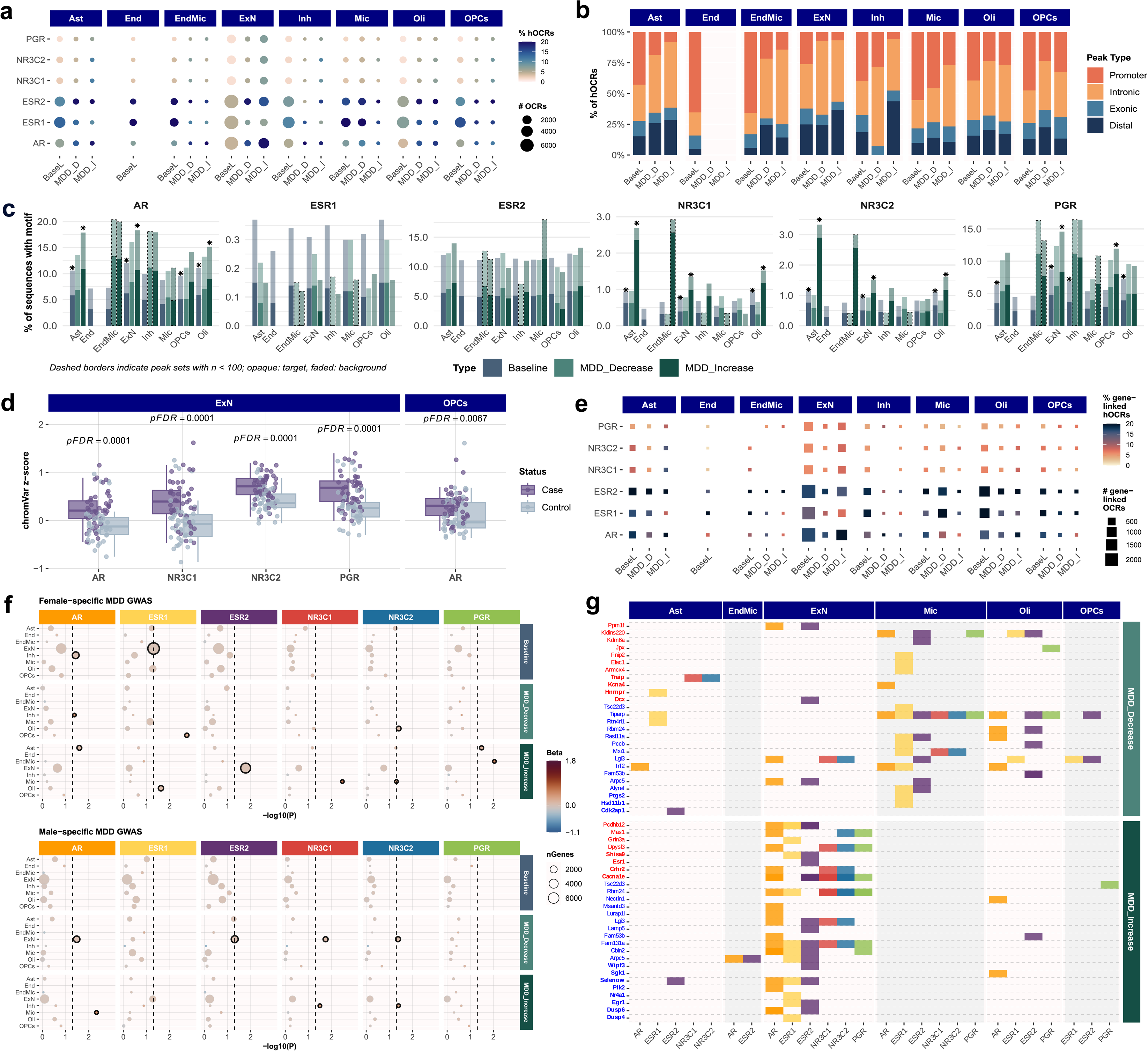
Hormone-responsive regulatory regions in depression show cell-type–specific motif enrichment and associate with sex-specific genetic vulnerability. (A) Dot plot showing the absolute count of OCRs per cell type per hormone motif per OCR category, coloured by proportion of hOCRs. (B) Stacked barplot showing the distribution of genomic context of hOCRs per cell type per OCR category. (C) Results from the HOMER enrichment analysis showing the enrichment of a given hormone motif in a given cell type per OCR category. Significant enrichments are indicated using an asterisk (pFDR<0.05). (D) Boxplots showing the distribution of chromVAR z-scores for hormone motifs in cases and controls. The chromVAR z-scores were aggregated to the subject level and only cell types that showed significant differences (pFDR<0.05) are shown. (E) Dot plot showing the absolute count of gene-linked OCRs per cell type per hormone motif per OCR category, coloured by proportion of gene-linked hOCRs. (F) Results from the MAGMA analysis showing sex-specific genetic liability for depression per cell type per hormone motif per OCR category. Significant enrichments are indicated in bold (p<0.05). (G) Overlap between hormone-linked genes and sex DEGs per cell type per depression-associated OCR category.

We next assessed motif enrichment to determine whether individual hormone receptor motifs show cell type–specific enrichment within each OCR category. HOMER^35^ analysis revealed that baseline OCRs were significantly enriched for *AR* and *NR3C1*/2 motifs in astrocytes, excitatory neurons, and oligodendrocytes, and for *PGR* motifs in excitatory neurons (Figure 5C). A similar pattern was observed for MDD_Increase OCRs, with additional *PGR* enrichment in OPCs, whereas MDD_Decrease OCRs showed no significant enrichment for hormone receptor motifs. These results suggest that gains in chromatin accessibility in depression are preferentially associated with hOCRs in select cell types, whereas decreases in accessibility show no consistent motif structure. Complementing this, chromVAR deviation scores^36^ showed that accessibility at hOCRs is significantly elevated in depressed cases in excitatory neurons (*AR*, *NR3C1/2* and *PGR*; pFDR=0.0001) and in OPCs for *AR* (pFDR=0.0067; Figure 5D; Figure S9), indicating that these hOCRs are not only overrepresented but are also actively more accessible in depression.

Having defined the hormone-responsive chromatin landscape, we next focused on gene-linked hOCRs to assess their biological relevance in depression. Gene-linked hOCRs showed patterns largely consistent with those observed across all hOCRs (Figure 5E; Figure S10), with a higher proportion of hOCRs across motifs, indicating widespread hormone-gene regulatory coupling. We then tested whether genes linked to these hOCRs are enriched for sex-specific depression polygenic risk using MAGMA^37^. At baseline, hormone-linked genes were enriched for female-specific depression heritability across *AR* and *ESR1/2*, particularly in excitatory and inhibitory neurons (p<0.05), with no corresponding enrichment in males (Figure 5F). This suggests that even within normative regulatory architecture, hormone-responsive gene networks may be more tightly coupled to inherited depression risk in females. In depression-associated OCRs, enrichment patterns were more context-dependent. Female-specific risk showed broader involvement of multiple cell types, including OPCs for *ESR1*, microglia for *NR3C1*, and endothelial–microglia for *PGR* (p<0.01). In contrast, male-specific enrichment was weaker and more restricted, with the strongest signal observed in microglia for *AR*, and additional nominal associations primarily in neuronal populations.

Finally, we examined the overlap between hormone-linked genes and DEGs identified in our dataset (Figure 5G; Figure S11). In total, 48 DEGs (including 19 depDEGs) mapped to depression-associated hOCRs. When stratified by cell type, DEGs linked to MDD_Increase OCRs were predominantly localized to excitatory neurons, whereas those linked to MDD_Decrease OCRs were primarily mapped to microglia. These patterns largely reflect the underlying hOCR landscape (Fisher’s test, pFDR>0.05), indicating no strong preferential targeting beyond baseline regulatory composition. Nevertheless, this analysis highlights candidate genes through which hormonal regulation may contribute to sex differences in depression. These include IEGs (*Nr4a1*, *Egr1*, *Dusp4/6*) linked to gonadal hOCRs in excitatory neurons within MDD_Increase regions, *Tiparp* (involved in aryl hydrocarbon receptor pathway) linked to all hOCRs in microglia within MDD_Decrease regions, and *Tsc22d3* which showed context- and cell type–specific associations with *ESR1* in microglia (MDD_Decrease) and *PGR* in excitatory neurons (MDD_Increase).

## DISCUSSION

Understanding sex differences in depression requires examining molecular processes across development rather than adult pathology alone. By integrating longitudinal multi-omic and gene regulatory network analyses in healthy animals, we identified normative baseline differences that may shape sex-specific responses to later stressors, consistent with a two-hit stress framework. Across analyses, these differences converge on three interconnected dimensions: divergent activity-dependent gene trajectories that shape the effects of pubertal stress, regulatory biases in stress-coping strategies, and hormone- and cell type–specific contexts through which genetic risk is expressed. Together, these findings position normative developmental regulation as a key substrate underlying sex-specific vulnerability to depression.

IEGs exhibited the strongest sex divergence in our study, with opposing developmental trajectories across adolescence (declining in females, increasing in males). IEGs function as activity-dependent markers of neuronal responsiveness and transcriptional inducibility, and in the context of existing literature, the direction of IEG expression changes may influence susceptibility to the long-term effects of pubertal stress, a known risk factor for depression. Gautier et al^38^ showed that although pubertal stress is associated with reduced baseline IEG expression in both sexes, subsequent exposure to allopregnanolone (AP; a progesterone-derived neurosteroid that rises during puberty, the luteal phase, and pregnancy) selectively induces IEG expression in females. This heightened inducibility may be due to increased chromatin accessibility established during the pubertal stress window, which may facilitate greater responsiveness to neuroendocrine feedback signals and contribute to the premature termination of the HPA response, a phenotype commonly observed following pubertal stress^39^. Integrating these findings with our data suggests that pubertal stress occurring during the downward trajectory of IEG expression in females may consolidate a suppressed yet more inducible baseline state. This may render the female HPA axis disproportionately sensitive to subsequent hormonal shifts, with AP acting as a sensitizer that triggers exaggerated IEG induction and aberrant downstream transcriptional responses. In contrast, males may be less vulnerable to this form of HPA dysregulation, as their higher adult IEG baseline may partially offset pubertal stress-induced suppression, alongside their ability to upregulate and utilize AP during acute stress for homeostatic recovery^40^. While it should be noted that the mechanistic evidence is derived from hypothalamic studies and may not be directly applicable to cortical systems, the opposing cortical IEG trajectories observed here raise the possibility that similar mechanisms may extend beyond the hypothalamus, warranting further investigation. Together, this framework of sex-specific IEG setpoints and divergent roles of AP may help explain why female depression risk rises sharply at puberty: a period when declining IEG baselines, rising AP, and stress exposure converge to create a more vulnerable HPA regulatory architecture.

In addition to divergent IEG trajectories contributing to differential vulnerability to depression, gene regulatory network analyses indicate that males and females differ in both the targets and control of stress-management systems. Under normative conditions, females exhibited tighter regulatory control over immune- and stress-related genes (e.g., *Fkbp5*, *Tsc22d3*), whereas males showed preferential regulation of activity-dependent and neuronal pathways (e.g., *Pink1*, *Camk2n1*). These patterns align with reported sex differences in stress coping, whereby males more effectively engage top-down prefrontal circuits during stress exposure, in contrast to females that showed reduced engagement of these circuits^41,42^, but greater GR reactivity during stress^43–45^. Here, we contextualize our normative findings in relation to stress using key genes as examples.

Male-biased regulatory architecture appears oriented toward maintaining neuronal stability. Evidence indicates that loss of *Pink1*, a mitochondrial kinase essential for mitophagy and also plays a role in modulating neuronal functions^46^, disproportionately affects males, who show earlier cognitive deficits^47^, increased depressive-like behavior, and lower hippocampal GR expression under chronic stress^48^. Maintaining higher baseline *Pink1* expression with activating upstream control, as observed here, may therefore be necessary for cognitive and stress resilience in males. Similarly, *Camk2n1* acts as a negative feedback regulator of CaMKII signaling^49^, whereby excessive excitation of CaMKII⁺ neurons during stress can initiate a pathological feedback loop that drives behavioral despair in male mice^50^. Maintaining higher baseline *Camk2n1* expression with repressive upstream control, as observed here, thus suggests a poised braking system that is constrained under normative conditions but available for rapid deployment to prevent stress-induced excitatory dysregulation. In contrast, female-biased regulatory architecture appears oriented toward prioritizing hormonal sensitivity. In Brygdes et al^43^, although males exhibited higher baseline expression of *Fkbp5*, glucocorticoid receptor (GR), and mineralocorticoid receptor (MR), prepubertal stress increased expression of *Fkbp5*, GR, and the GR:MR ratio only in females, suggesting that the system is actively adapting to stress. Notably, only prepubertally stressed females showed a blunted HPA response during adult social stress testing. Relating these to our findings, it is likely that females exhibit lower baseline expression but tighter, more balanced transcriptional control of *Fkbp5* to preserve GR sensitivity and effective HPA-axis feedback during stress. In parallel, the lower baseline expression of *Tsc22d3* coupled with activating upstream regulation suggest a system primed for rapid induction upon GR activation. These features are consistent with a regulatory architecture tuned for heightened GR responsiveness and immune engagement in females.

Taken together, these findings suggest that sex-specific regulatory architectures supporting neuronal stability in males and hormonal sensitivity in females, while adaptive under normative conditions, may become liabilities when regulatory control is compromised. In this context, males may be more vulnerable to stress-induced circuit dysregulation, whereas females may be more susceptible to amplified GR signaling and impaired HPA-axis feedback. These differences define distinct entry points for stress-induced pathology and provide a mechanistic basis for how comparable stress exposures can lead to divergent depression risk and outcomes across sexes.

The final dimension concerns the differential involvement of hormone-responsive cell-type–specific contexts in depression. Our analyses highlight preferential enrichment of specific cell types, such as astrocytes and excitatory neurons, across both sexes, indicating that these populations may play a disproportionate role in depression-associated regulatory changes^51–53^. Furthermore, we find that hormone-responsive regulatory programs shape the translation of genetic risk into cell-type–specific transcriptional dysregulation in a sex-specific manner. Mechanistically, this may occur through risk variants altering TF binding motifs and thereby disrupting regulatory activity, as demonstrated in depression-relevant contexts^34^. Although functional validation of the observed genetic enrichment remains necessary, existing evidence supports a role for hormone–cell type interactions in stress- and depression-related transcriptional regulation.

For example, female-specific genetic liability for depression is enriched in *PGR*-linked endothelial–microglia clusters (key players at the BBB), consistent with evidence of sex-dependent vascular vulnerability in stress-related pathology^54–56^. Chronic social stress induces loss of the tight-junction protein *Cldn5* and increased BBB permeability selectively in females, with substantial baseline sex differences in endothelial gene expression already present in unstressed controls^54,56^, suggesting that female vulnerability may arise from a pre-configured vascular regulatory landscape that is amplified under stress. Further, although progesterone is often neuroprotective, this effect is inconsistent in females^57,58^, suggesting reduced engagement of protective progesterone-responsive programs at the vascular-immune interface, in line with the divergent roles of AP discussed above. This enrichment was observed in MDD_Increase OCRs, consistent with preferential engagement of stress- and immune-related regulatory programs in female depression. Conversely, the enrichment in *Esr1*-linked OPCs within MDD_Decrease OCRs suggests that reduced ERα-mediated OPC differentiation, remyelination, and antidepressant effects during stress^59^ may contribute towards vulnerability.

On the other hand, male-specific genetic risk converged on androgen- and glucocorticoid-responsive neuroimmune and neuronal contexts, respectively. Enrichment in *AR*-linked microglia clusters within MDD_Increase OCRs is consistent with evidence that androgen signaling contributes to neuroimmune regulation and microglial *AR* expression may be governed by distinct regulatory mechanisms in males^60^. Imbalance between androgen and estrogen signaling can also disrupt microglial homeostasis and contribute to depressive-like behaviours males^61^. Enrichment in *NR3C1/2*-linked neuronal clusters within both MDD_Increase and MDD_Decrease OCRs is consistent with the role of corticosteroids in regulating NMDA-receptor signaling in hippocampal neurons^62^, and mediating stress-related behavioural responses through regulation of key neurotransmitter systems^63^. This suggests that deviations in glucocorticoid-mediated neuronal regulation, in either direction of chromatin accessibility change, are associated with MDD-related regulatory states in males. Together, these results indicate that immune- and glucocorticoid-responsive programs are also engaged in males but are expressed through distinct hormonal and cellular contexts. In this case, they appear preferentially embedded within androgen-dominant signaling environments, reflecting the dominant hormonal milieu in males, and within neuron-centric regulatory states, reflecting a male-biased regulatory strategy oriented toward neuronal stability. These findings underscore the importance of cell-type–resolved analyses for interpreting sex differences in stress-related regulatory processes.

Collectively, these findings support a developmental framework in which sex differences in depression vulnerability and outcomes emerge from divergent activity-dependent gene trajectories that shape responses to pubertal stress, sex-specific regulatory architectures governing stress responses, and hormone- and cell type–specific contexts through which genetic risk is expressed. These processes unfold across critical developmental periods, highlighting the importance of early regulatory architecture in shaping later vulnerability, such that disruption during sensitive windows may lead to persistent dysregulation along distinct biological and clinical trajectories. This provides a basis for how shared risk factors give rise to heterogeneous outcomes across sexes, including differences in prevalence, symptom presentation, and treatment response.

A major strength of this study is the high temporal resolution with which sex-specific gene regulatory dynamics were modelled, allowing us to capture dynamic changes in gene expression and regulatory architecture. Integrating multiple analytical layers also allowed us to interrogate these processes from complementary regulatory perspectives. Despite these strengths, several limitations should be considered when interpreting our findings. First, all analyses were conducted in bulk RNA-seq data from the prelimbic cortex. The sex-specific regulatory strategies identified here may not generalize to other brain areas implicated in depression, nor do they capture cell-type–resolved effects within the cortex. Second, this study does not directly model depressive pathology. While this was intentional to characterize normative sex differences and potential vulnerabilities, it limits inference about disease causality. Third, our developmental window began at P28, but substantial brain and hormonal programming occurs earlier in life^64^, which may shape trajectories in ways not captured here.

These limitations suggest several avenues for future work. Experimental validation will be essential to test the functional relevance of the identified regulatory strategies. Targeted perturbation of key genes (e.g., *Fkbp5* in females, *Camk2n1* in males) during adolescence could help determine whether disrupting these buffering mechanisms increases susceptibility to depression-like behaviors. Extending these analyses to single-nucleus data from postmortem human brain tissues, together with direct measures of brain steroid hormones, would further resolve cell-type–specific mechanisms and assess the translational relevance of the observed sex-specific hormone-sensitive regulatory architectures.

## MATERIALS AND METHODS

### Animals

Wild-type C57BL/6NCrl mice were obtained from the Charles River Laboratories (Laval, Québec, Canada) at postnatal day 21 (P21). The mice were habituated to the animal facility for one week prior to experimentation and housed under standard conditions (22°C, 12h light/dark cycle) with ad libitum access to food and water. Animals were housed in groups with same-sex littermates. Transcriptomic and methylation data were obtained from independent cohorts of animals. For transcriptomic data, two animals per sex were euthanized by swift decapitation without anaesthesia at each of the 21 time points (P28, P32, P36, P38, P40, P42, P44, P46, P48, P50, P52, P54, P56, P58, P60, P62, P64, P68, P70, P72, and P120). For methylation data, three animals per sex were euthanized at P28, followed by three males and six females for each of the four subsequent time points (P38, P42, P48, and P72). All animals were euthanized during the dark phase between 8:30 and 9:30pm. All procedures were conducted in accordance with the guidelines of the Canadian Council of Animal Care and the McGill University/Douglas Mental Health University Institute Animal Care Committee.

### Tissue collection and microdissection

Following rapid decapitation, whole brains were immediately extracted and snap-frozen in isopentane placed on dry ice to preserve tissue integrity. Frozen brains were stored at −80°C until further processing. Coronal sections (200 μm-thick) were prepared on a Leica CM1950 Cryostat (Leica Biosystems, Deer Park, IL) maintained at −16°C. Sections were mounted onto Fisherbrand™ Superfrost™ Plus microscope slides and kept on dry ice during dissection to maintain tissue stability. Bilateral punches of the region of interest (0.75 mm diameter) were collected and stored at -80°C until nucleic acid extraction.

### Transcriptomic and DNA methylation profiling

For the transcriptomic dataset, total RNA was extracted from the prelimbic subregion of the medial prefrontal cortex using the RNeasy Lipid Tissue Mini Kit (Qiagen, Cat #74804). On-column DNase digestion was performed with the RNase-Free DNase Set (Qiagen) to eliminate DNA contamination. Bulk mRNA libraries were constructed using the Illumina’s Stranded mRNA Prep Kit (Cat #20040534) according to the manufacturer’s protocol. The libraries were pooled and sequenced at Genome Québec on the Illumina NovaSeq platform. For the methylation dataset, DNA was extracted from the entire medial prefrontal cortex (mPFC) using the QIAamp DNA Micro Kit (Qiagen, Cat #56304). Bisulfite conversion was performed at Genome Québec using the EZ-96 DNA Methylation-Gold™ Kit (Zymo Research, Cat #D5008). Bulk DNA methylation profiling was conducted by hybridizing bisulfite-converted DNA to the Infinium Mouse Methylation BeadChip (Illumina, Cat #20041558), a high-resolution array covering 285,000 CpG sites across the mouse genome. Array processing and scanning were conducted according to the manufacturer’s protocol.

### Data preprocessing

For the transcriptomic dataset, genes with fewer than 10 counts in at least 10 samples were removed. Counts were normalized for library size and variance-stabilized using the regularized log (rlog) transformation in DESeq2^65^. To improve translational relevance, only genes with human orthologs were retained for downstream analyses, resulting in 15,177 genes across autosomes and chromosome X. For the methylation dataset, raw Illumina array data were preprocessed using the meffil^66^ pipeline, which included sample quality control, probe filtering, and quantile normalization of beta values. Poor-quality probes identified during QC were removed, and normalized methylation values were generated following standard meffil procedures. CpGs were annotated to genes using the Illumina annotation file, and only CpGs mapping to genes with human orthologs were retained for downstream analyses, yielding 154,651 CpGs across autosomes and chromosome X.

### Cell-type deconvolution

Reference-based cell type deconvolution was done to estimate the relative contribution of individual cell types in the bulk transcriptomic and methylation data. The single-cell RNA-seq atlas^67^ and the single-cell methylome atlas^68^ of the mouse brain were used as references for the transcriptomic and methylation datasets, respectively. Deconvolution was implemented using CIBERSORTx^69^, and principal component analysis (PCA) was done across cell types to derive principal components. We then tested whether proportions of each cell type differed significantly (1) between sexes and (2) across ages using linear regression. If any cell type exhibits a significant difference, PCA was done and PC1 was included as a covariate in downstream statistical models to ensure that observed effects were not confounded by differences in that cell type.

### Broad developmental analyses including adolescence and adulthood

#### Generating and characterizing iDREM gene regulatory networks

iDREM is a computational framework that integrates time-series gene expression data with known or predicted regulatory information (e.g., transcription factor (TF)-motif, DNA methylation) to construct dynamic gene regulatory networks (GRNs)^30^. Briefly, it utilizes Hidden Markov Models to (1) map gene expression trajectories, (2) identify bifurcation points where co-expressed genes begin to diverge in their expression patterns, and (3) infer the regulators most likely driving these divergences. In this study, iDREM was used to determine when and where key regulatory events occur from P28 to P120. Analyses were performed separately for each sex, using our time-series transcriptomic and methylation data, along with curated TF-motif information from the iDREM database (https://github.com/phoenixding/idrem/tree/master/TFInput). iDREM compares the average expression of the genes within each path to their expression at baseline (P28; designated as nodeMean=0) – paths with expression levels higher than zero represent an upregulation of expression relative to the baseline, vice versa. Each gene has exclusive membership to a specific path. Here, we focused mainly on genes belonging to paths that were upregulated or downregulated relative to baseline in each sex. Upregulated paths were defined as those in which at least 50% of nodes had a nodeMean>0, and downregulated paths were defined as those in which at least 50% of nodes had a nodeMean<0. Pathway enrichment analysis of FUMD (female up, male down) and FDMU (female down, male up) genes was conducted using the *msigdbr* package across diverse gene set collections, such as GO, Reactome, HP, PID, and KEGG. To identify regulators involved in driving each path, we looked at the bifurcation nodes (they represent the initiators of a path) and end nodes (they represent the maintainers of the path). For interpretation, only the top 3 regulators of each path in each category based on their p-values were extracted.

#### Transcriptomic maturation analysis

Transcriptomic maturation was quantified using a distance-based framework adapted from Kaplan et al^64^. We computed the distance in PC space between each pre-adult timepoint and the adult reference (P120), separately for three gene sets: global (using the top 2000 highly variable genes), FUMD, and FDMU genes. For each gene set, expression matrices were z-score normalized per gene and PCA was conducted jointly across both sexes to define a shared transcriptional space. The number of PCs retained was defined as the minimum required to explain ≥80% of the total variance. Within this shared PC space, for each sex, age-specific centroids were derived and Manhattan distance between each centroid and the P120 centroid was calculated. To ensure the trajectory reflected cumulative maturation rather than transient fluctuations, a monotonicity constraint was applied: where distance increased between consecutive timepoints, the later value was replaced by the preceding one.

### Statistical modelling of adolescent molecular dynamics

Statistical modeling of adolescence was restricted to timepoints P28 to P72, as the P120 timepoint is distal and may introduce bias in model estimates. We implemented a unified analytical framework based on generalized additive models^70^ (GAMs) and correlation-based integration to characterize sex-specific regulatory dynamics across gene expression, DNA methylation, and TF regulatory networks.

#### Gene expression analyses

To assess whether gene expression differs between sexes across time, mass univariate testing was performed for each gene using GAMs with a cubic smoothing function (*mgcv* R package): *gam(expression ∼ sex + s(age, by = sex)).* False discovery rate (FDR) correction was applied across all genes, with significance defined at pFDR<0.05. Genes significant for the main sex term were interpreted as having differences in average expression between sexes. To identify genes with sex-dependent expression trajectories, an ANOVA was performed by comparing the full model to a reduced model (*gam(expression ∼ sex + s(age))*). Differentially expressed genes (DEGs) were defined as genes with a significant main effect of sex and/or a significant sex-by-age interaction.

To identify DEG clusters, predicted expression trajectories were generated for each DEG using 100 evenly spaced age points between P28 and P72. Trajectories were scaled to the range [0,1] and hierarchically clustered using Ward’s linkage with Euclidean distance. Consensus clustering was applied to evaluate a range of cluster solutions. Based on inspection of the consensus heatmap and dendrogram structure, the hierarchy was cut to yield four clusters, capturing the predominant expression patterns while minimizing redundancy. Pathway enrichment analysis for each DEG cluster was performed using gene sets from the *msigdbr* package.

Depression-associated genes were curated from three sources: the most recent depression genome-wide association study (GWAS) by the Psychiatric Genetics Consortium^71^, the DISGENET database (focusing on mental depression; CUI: C0011570) and major depressive disorder (CUI: C1269683), and a single-nucleus RNA-seq study of postmortem dorsolateral prefrontal cortex tissue from individuals with depression^72^. Only genes with mouse orthologs were retained, resulting in a curated list of 2580 depression-associated genes.

#### DNA methylation analyses

The same GAM framework was applied to assess whether DNA methylation levels differed between sexes across time: *gam(methylation ∼ sex + s(age, by = sex))*. FDR correction was applied across all CpGs, with significance defined at pFDR<0.05. Differentially methylated positions (DMPs) were defined as CpGs exhibiting a significant main effect of sex and/or a significant sex-by-age effect on methylation trajectories. CpGs were annotated using the *annotatr* R package, assigning genomic features according to a hierarchical priority (e.g., promoter, exon, intron, intergenic). Differentially methylated regions (DMRs) were defined by grouping DMPs within a maximum gap of 1000 bp, requiring at least three DMPs per region. Hormone motifs for *Ar*, *Esr1*, *Esr2*, *Nr3c1*, *Nr3c2* and *Pgr* were identified using the *JASPAR2020* R package. Overlaps between DMRs and hormone motifs were identified using the *motifr* package with a significance threshold of p<0.0001.

To assess relationships between methylation and gene expression dynamics, predicted methylation trajectories were generated for each CpG within a DMR using 100 evenly spaced age points between P28 and P72. The mean predicted methylation trajectory across CpGs in a DMR was then correlated with the corresponding predicted gene expression trajectory using a hybrid metric combining distance and Spearman correlation. Distance correlation quantifies the overall strength of association between variables, capturing both linear and non-linear dependencies (ranging from 0, indicating independence, to 1, indicating strong dependence), whereas Spearman correlation assesses monotonic relationships and provides directional information. As distance correlation is non-directional, a signed metric was computed by multiplying the distance correlation magnitude by the sign of the Spearman correlation. Statistical significance was assessed using block permutation (n=10,000). For each permutation, expression trajectories were divided into four contiguous blocks (best reflecting developmental epochs) and randomly shuffled to generate a null trajectory, which was then correlated with the methylation trajectory. This approach preserves local temporal structure within blocks while disrupting global alignment between trajectories, providing a null model that accounts for autocorrelation and aligns with block-based resampling strategies for dependent data^73^. Empirical p-values were computed as the proportion of null correlations exceeding the observed value. As this analysis was intended to identify candidate associations within a structured set of temporal trajectories, FDR correction was not applied. Associations were instead evaluated using nominal significance (p<0.05) for either distance or Spearman correlation, reflecting complementary sensitivity to non-linear and monotonic relationships. The identified associations were mainly of large effect size (|r|>0.5), supporting the robustness of the observed relationships.

#### TF targeting analyses

PANDA is conceptually similar to iDREM in that it integrates multiple sources of evidence to infer gene regulatory relationships^74^. It employs a message-passing framework to iteratively update edge weights between TFs and target genes, yielding a group-level weighted bipartite network that reflects inferred regulatory influence under a given condition (e.g., sex). Edge weights indicate the strength of evidence for a regulatory relationship, with strong positive values representing stronger evidence and zero or negative values indicating no or weak evidence. LIONESS extends PANDA by enabling reconstruction of sample-specific GRNs from the group-level network using a leave-one-out approach, thereby estimating each sample’s contribution to the overall regulatory structure^33^. Analyses were performed separately for each sex using our time-series transcriptomic data, TF motif information from the iDREM database, and protein–protein interaction data from STRINGdb. Each raw sample-specific GRN contained approximately 4.6 million edges. To reduce noise and focus on robust interactions, edges were filtered to retain those with edge weights > 1 in at least 50% of samples within each sex, and only edges involving genes with human orthologs were retained, resulting in 871,037 edges.

To assess whether TF-gene regulatory interaction strength differs between sexes across time, the same GAM framework described above was applied to all edge weights: *gam(edge_weight ∼ sex + s(age, by = sex))*. FDR correction was applied across all edges, with significance defined at pFDR<0.05. Sex-biased regulatory edges were defined as those exhibiting a significant main effect of sex and/or a significant sex-by-age effect. To quantify associations between TF regulatory trajectories and gene expression trajectories, the same permutation-based framework described above was applied to the 1134 edges involving DEGs. Associations were evaluated using a relaxed nominal threshold (p<0.1) for either distance or Spearman correlation to prioritize candidate regulatory relationships for downstream interpretation. The identified associations were predominantly of large effect size (|r|>0.5), supporting the robustness of the observed relationships. As edge weights are not directional, the sign of the correlation was used to indicate whether changes in regulatory interaction strength were positively or negatively associated with gene expression, consistent with putative activating or repressing relationships. To summarize regulatory directionality across TFs for each DEG, a balance ratio was computed as the proportion of putatively activating edges relative to the total number of edges associated with each TF. To identify TFs exhibiting sex-specific regulatory patterns, edges were first classified as shared (significant in both sexes), male-specific, or female-specific (significant in only one sex). For shared edges, regulatory directionality was defined by the sign of the correlation coefficient, and consistency between sexes was assessed based on sign concordance. The proportion of concordant and discordant shared edges for each TF was derived, and the average correlation strength for shared edges within each sex was calculated using Fisher’s z-transformation.

#### Hormone-motif analyses

Analyses were performed using single-nucleus ATAC-seq data derived from adult human postmortem dorsolateral prefrontal cortex^34^. The original study focused on differential chromatin accessibility between major depressive disorder (MDD) cases and controls. Here, we analyzed three categories of open chromatin regions (OCRs): baseline OCRs (N=336,728 after selecting for highly reproducible peaks), MDD_Increase OCRs (N=19,363), and MDD_Decrease OCRs (N=14,452). In the absence of direct hormone measurements, a motif-based approach was used to infer potential hormone responsiveness, whereby the presence of a hormone response motif within an OCR suggests putative regulatory potential. Position weight matrix files for hormone motifs were obtained from the JASPAR database, including *AR* (MA0007.2), *ESR1* (MA0112.3), *ESR2* (MA0258.2), *NR3C1* (MA0113.3), *NR3C2* (MA0727.2), and *PGR* (MA2327.1). Motif scanning was performed using FIMO (Find Individual Motif Occurrences)^75^ with a significance threshold of p<0.0001. In this context, the FIMO p-value reflects the likelihood of observing a motif match of equal or greater similarity to the position weight matrix by chance within a given sequence. Hormone-responsive OCRs (hOCRs) were then annotated to genes using the peak-to-gene (p2g) linkage file derived previously^34^.

Cell-type-specific hormone motif enrichment was assessed using HOMER (Hypergeometric Optimization of Motif EnRichment)^35^. For each cell type, OCRs containing hormone motifs were tested against a background of all OCRs from the remaining cell types. FDR correction was applied across tests, with significance defined at pFDR<0.05. For motif accessibility analysis, GC content and mean accessibility bias-corrected motif accessibility deviation *z*-scores were computed at the single-cell level using chromVAR^36^ (v.1.14.0) based on the Cisbp motif database in the addDeviationsMatrix() function in ArchR. The chromVAR z-scores per cell type were averaged to the subject level and the Wilcoxon signed-rank test was used to test for case-control differences. FDR correction was applied across tests, with significance defined at pFDR<0.05.

Gene-level enrichment analyses were performed using MAGMA (Multi-marker Analysis of GenoMic Annotation)^37^, leveraging summary statistics from a sex-specific depression GWAS^76^. To meet the minimum gene set size requirements for reliable MAGMA analysis, a more permissive p2g correlation threshold (>0.2) was used in conjunction with pFDR<0.05 to map hOCRs to genes. To ensure that enrichment of MDD GWAS signal in hormone-regulated OCRs reflected hormone specificity rather than general cell-type-level GWAS enrichment, we conditioned each gene set on a background of all OCR-linked genes in the same cell type. For baseline hormone-regulated OCRs, the background was all genes linked to OCRs in that cell type regardless of hormone motif presence. For depression-associated hormone-regulated OCRs, the background was all genes linked to depression-associated OCRs in that cell type. This conditional analysis therefore asks whether hormone-regulated genes show enrichment above what would be expected given the cell type’s overall GWAS signal. Due to the exploratory nature of this analysis and the lack of statistical power from the limited number of hOCRs, FDR correction was not applied to MAGMA results and nominal p-values were reported.

We used a more stringent p2g linkage threshold (correlation>0.4 and pFDR<0.05) to map hOCRs to genes before overlapping the list of genes with sex DEGs. To test whether sex DEGs were enriched among hormone target genes within depression-associated OCR-linked genes, we performed hypergeometric enrichment analyses stratified by cell type and OCR category. For each test, the gene universe was defined as all genes linked to OCRs in the corresponding cell type. Within this universe, we tested whether the overlap between sexDEGs and hormone target genes was greater than expected by chance under a hypergeometric model. FDR correction was applied across tests, with significance defined at pFDR<0.05.

## Author contributions

G.T conceptualized the study, performed the analyses, interpreted the results, generated the figures, and wrote the manuscript. M.G. generated data, contributed to the development of the analytical framework, and documentation of the study. K.S and J.D performed iDREM and TPS analyses. A.C provided the single-nucleus ATAC-seq data and conducted chromVAR analyses. M.S processed the DNA methylation data. R.K, D.L, T.Y.Z and T.P.W contributed to data generation. M.N and G.F provided bioinformatics support. N.S and Y.Z provided methodological expertise. C.N conceptualized the study, oversaw the project, contributed to manuscript development and editing, and provided funding. All authors reviewed the manuscript.

## Acknowledgements

G.T is supported by the National Science Scholarship from the Agency of Science, Technology and Research, Singapore. C.N. is supported by grants from NSERC (RGPIN-2022-03979), NARSAD (BBRF #31204) and CIHR (PJT183904).

## Code availability

All code used in this study will be available at https://github.com/MGSSdouglas/LongitudinalMouseGRN

